# TbasCO: Trait-based Comparative ’Omics Identifies Ecosystem-Level and Niche-Differentiating Adaptations of an Engineered Microbiome

**DOI:** 10.1101/2021.12.04.471239

**Authors:** E.A. McDaniel, J.J.M van Steenbrugge, D.R. Noguera, K.D. McMahon, J.M. Raaijmakers, M.H. Medema, B.O. Oyserman

**Author notes:** corresponding author Corresponding authors: Elizabeth McDaniel, Joris van Steenbrugge, Ben Oyserman. contributed equally (E.A. McDaniel, J.J.M van Steenbrugge). Department of Microbiology and Immunology, University of British Columbia, Vancouver, CA.

## Abstract

A grand challenge in microbial ecology is disentangling the traits of individual populations within complex communities. Various cultivation-independent approaches have been used to infer traits based on the presence of marker genes. However, marker genes are not linked to traits with complete fidelity, nor do they capture important attributes, such as the timing of expression or coordination among traits. To address this, we present an approach for assessing the trait landscape of microbial communities by statistically defining a trait attribute as shared transcriptional pattern across multiple organisms. Leveraging the KEGG pathway database as a trait library and the Enhanced Biological Phosphorus Removal (EBPR) model microbial ecosystem, we demonstrate that a majority (65%) of traits present in 10 or more genomes have niche-differentiating expression attributes. For example, while 14 genomes containing the high-affinity phosphorus transporter *pstABCS* display a canonical attribute (e.g. up-regulation under phosphorus starvation), we identified another attribute shared by 11 genomes where transcription was highest under high phosphorus conditions. Taken together, we provide a novel framework for revealing hidden metabolic versatility when investigating genomic data alone by assigning trait-attributes through genome-resolved time-series metatranscriptomics.

## INTRODUCTION

A longstanding cornerstone of deterministic ecological theory is that the environment selects for traits. Traits may be defined as any physiological, morphological, or genomic signature that affects the fitness or function of an individual [1]. Trait-based approaches have become indispensable in macroecological systems to describe fitness trade-offs and the effects of biodiversity on ecosystem functioning [2–5]. Recently, trait-based frameworks have been proposed as an alternative to taxonomy-based methods for describing microbial ecosystem processes [6, 7]. Connecting microbial traits and their phylogenetic distributions to ecosystem performance can provide powerful insights into the ecological and evolutionary dynamics underpinning community assembly, microbial biogeography, and organismal responses to changes in the environment [8–10]. Additionally, pinpointing the organismal distribution of traits and the selective pressures that enrich them may be leveraged to reproducibly and rationally engineer stable, functionally redundant ecosystems [11–15]. However, applying trait-based approaches to microbial communities is challenging due to the difficulty in identifying and measuring relevant ecological traits for a given ecosystem [16].

High-throughput sequencing technologies and multi-omics techniques have been used to describe the diversity, activity, and functional potential of uncultivated microbial lineages [17–20]. Improvements in bioinformatics algorithms, and in particular metagenomic binning methods, have allowed for genome-resolved investigations of microbial communities rather than gene-based analyses of assembled contigs [21]. These (meta) genomes are subsequently leveraged to detect the presence of key genes or pathways and predict specific traits of the whole community [22, 23]. Integrating metatranscriptomics data addresses a key limitation, as expression patterns better reflect the actual functional dynamics of a trait compared to gene presence alone. Here, we present TbasCO, a software package and statistical framework for *T*rait-*bas*ed *C*omparative ‘*O*mics to identify expression attributes. We adopt the terminology *attribute* as a hierarchically structured feature of a trait and assert that statistically similar transcriptional patterns of traits across multiple organisms be treated as *attributes* (Figure 1). In this manner, the identification of expression-based *attributes* provides a high-throughput and intuitive framework for extending trait-based methods to time-series expression patterns in microbial communities. We implement this trait-based approach to classify transcriptional attributes in a microbial community performing Enhanced Biological Phosphorus Removal (EBPR), a globally important biotechnological process implemented in numerous wastewater treatment plants (WWTPs).

**Figure 1.**
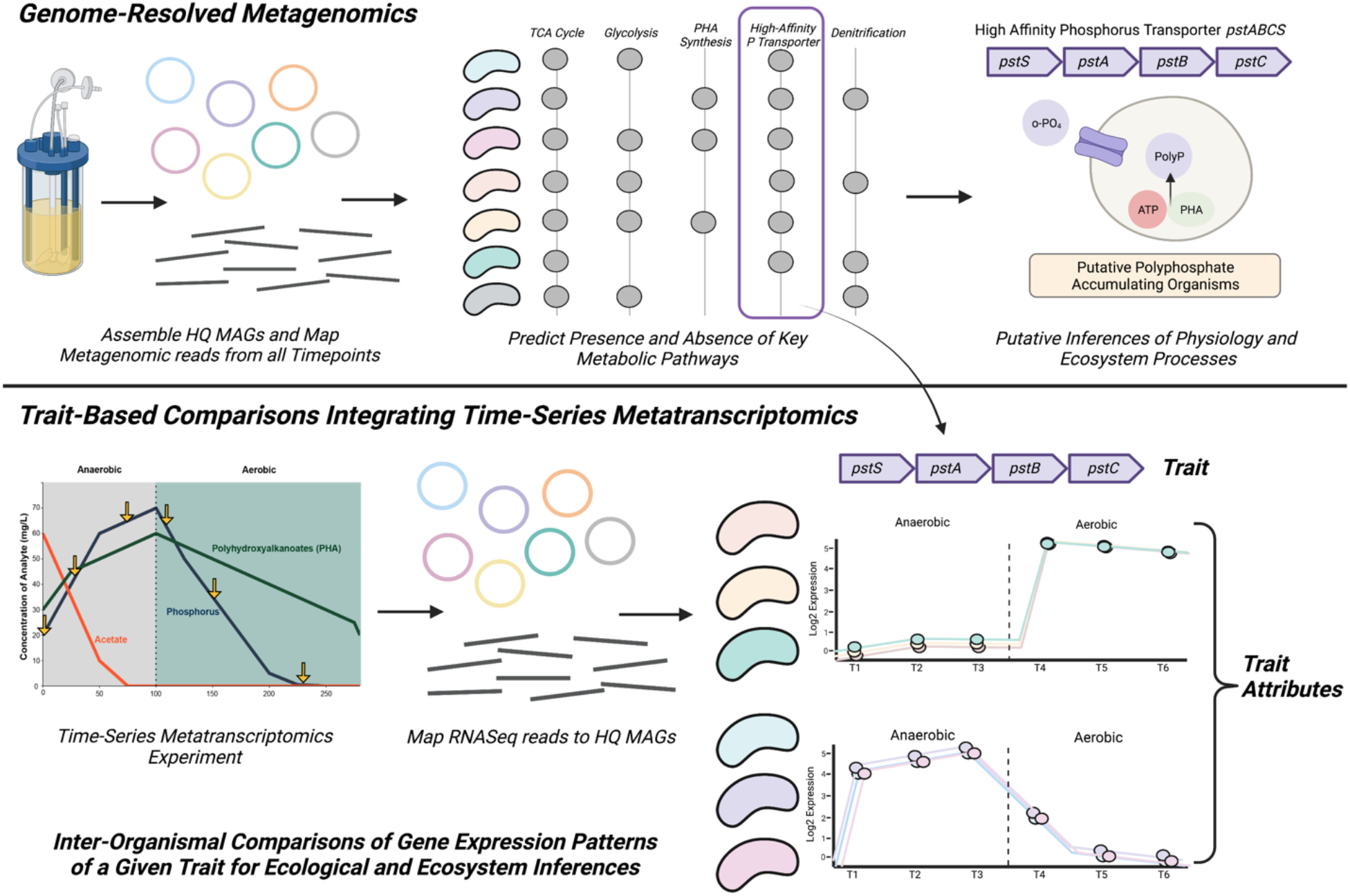
Overview of Trait-based Comparative Transcriptomics Approach. In genome-resolved metagenomics approaches, representative MAGs are assembled from a microbial community of interest, and the presence and/or absence of key metabolic pathways are used to make inferences of metabolic potential and ecosystem processes. However, metagenomic data alone can only assess the metabolic potential of a given pathway, and do not provide other biologically relevant information such as the timing or induction of these traits. Using time-series metatranscriptomics, we developed a trait-based comparative ‘omics (TbasCO) pipeline that statistically assesses the inter-organismal differences in gene expression pattern of a given trait to cluster into trait attributes.

The fundamental feature of the engineered EBPR ecosystem is the decoupled and cyclic availability of an external carbon source and terminal electron acceptor. This cycling is often referred to as “feast-famine” conditions and provides a strong selective pressure for traits such as polymer cycling. Accumulation of intracellular polyphosphate through cyclic anaerobic-aerobic conditions ultimately results in net phosphorus removal and accomplishes the EBPR process [24, 25]. One of the most well-studied polyphosphate accumulating organisms (PAOs) belongs to the uncultivated bacterial lineage ‘*Candidatus* Accumulibacter phosphatis’ (hereby referred to as Accumulibacter) [24, 26]. Numerous genome-resolved ‘omics methods have been used to investigate the physiology and regulation of this model PAO enriched in engineered lab-scale enrichment bioreactor systems [27–34]. However, novel and putative PAOs have been discovered that remove phosphorus without exhibiting the hallmark traits of Accumulibacter [35–39]. Additionally, although these lab-scale systems are designed to specifically enrich for Accumulibacter, a diverse “flanking community” persists in these environments [27], and their ecological roles have largely remained unexplored. As a result, the general adaptations of microbial lineages inhabiting the EBPR community are not well understood. Using genome-resolved metagenomics and metatranscriptomics, we assembled 66 species-representative genomes spanning several significant EBPR lineages and identified the distribution of expression-based attributes. Using our novel trait-based comparative ‘omics approach, we show that while some expression attributes are distributed in few genomes, many are redundant and shared across many lineages. Furthermore, we find that a majority of core traits (as defined by the presence of marker genes) have multiple attributes, suggesting that identifying niche-differentiating expression attributes may be used to reveal a large hidden metabolic versatility when investigating genomic data alone.

## MATERIALS AND METHODS

### Metagenomic Assembly, Annotation, and Metatranscriptomic Mapping

Three metagenomes sampled from an EBPR bioreactor with linked time-series metatranscriptomics data [40] were collected for metagenomic sequencing and assembled into 66 species-representative bins as described in the Supplemental Methods. All bins are greater than 75% complete and contain less than 10% contamination, with a large majority (44/66) >95% complete and <5% redundant as calculated by CheckM [41] (Table 1). Each bin was functionally annotated using the KEGG database through an HMM-based approach under KEGG release 93.0 using the command-line KofamKOALA pipeline [42, 43], selecting annotations that were significant hits above the specific HMM threshold. This resulted in 117,657 total annotations with 5,228 unique annotations. We used a metatranscriptomic dataset of six timepoints collected over a single EBPR cycle from Oyserman et al. 2016 [40], with three timepoints from the anaerobic phase and three from the aerobic phase. Raw reads were quality filtered using BBtools suite v38.07 [44] and ribosomal rRNA was removed from each sample using SortMeRNA [45]. Reads from each sample were mapped against the concatenated set of open reading frames from all 66 bins using kallisto v0.44.0 and parsed using the R package tximport [46, 47].

**Table 1.**
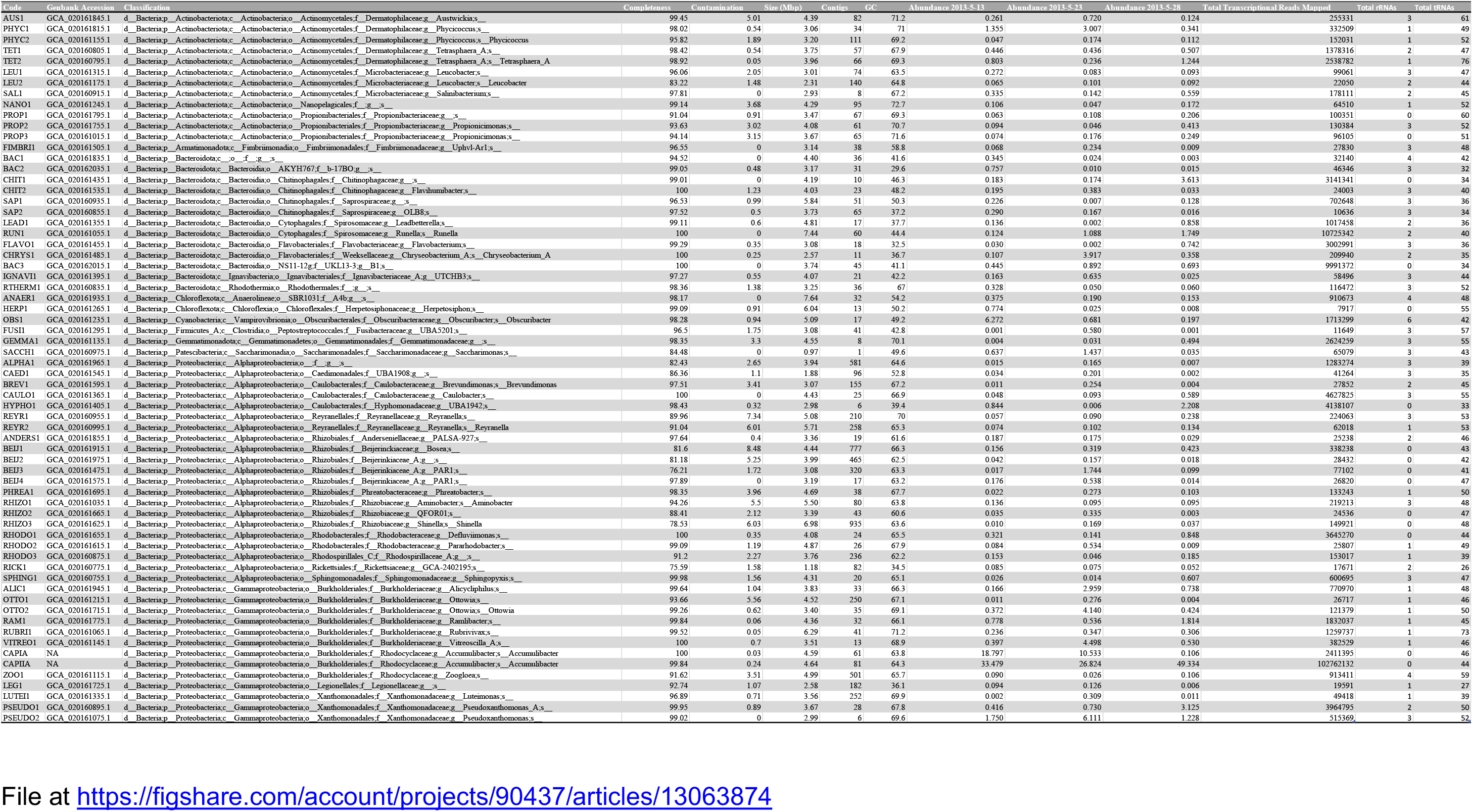
Genome quality statistics and relative abundance calculations for all 66 EBPR SBR MAGs. Genome code names match names used in all figures and within the text. Classifications were assigned using the GTDB-tk [87] and confirmed by comparing against select publicly available references and a subset of HQ MAGs from Singleton et al. 2021 [39]. Completeness and redundancy estimates and GC content were calculated by CheckM [41]. tRNA and rRNA predictions were performed with Barrnap as part of the Prokka software [88]. Relative abundance estimates reflect the proportion of reads mapped to the genome in that sample divided by the total number of reads mapped to all genomes as performed with SingleM. Table available at https://figshare.com/articles/dataset/EBPR_SBR_MAGs_Metadata/13063874.

### TbasCO Method Implementation

The TbasCO package identifies expression-based attributes of predefined traits using time-series (meta)transcriptomics data (Figure 1). In general, traits are defined by the presence of a pathway or other collection of genes from an externally provided database. A weighted distance metric between expression patterns for all genes that define a trait is calculated, and statistically significant similarity is determined based on the background distribution of a trait of equal size. Thereby, two or more organisms with a statistically similar expression pattern for a trait share an *attribute*.

### Input and Preprocessing

The input that is accepted by TbasCO is a table of RNAseq counts in csv format. Each row is treated as gene that has columns for the gene/locus name, counts per sample, the genome the gene belongs to, and the KEGG Orthology (KO) identifier. The RNAseq counts table may be provided pre-normalized or can be normalized by the program. The default normalization method is designed to minimize compositional bias in the differential abundance and activity of constituent populations in metatranscriptomics studies. Raw RNA expression counts are therefore normalized by genomic bin and sample [40]. These normalization factors are then applied to each sample for each bin individually. Alternatively, custom normalization methods may be implemented. After normalization, a pruning step is introduced to filter genes that have zero counts or a mean absolute deviation of less than one. To make inter-organismal comparisons of the relative contribution of a gene to total measured organismal RNA, an additional statistic is calculated ranking the expression counts from each sample from highest to lowest. The ranks for each sample are then normalized by dividing them by the maximum rank value in that sample. This normalization is applied to make ranks comparable between organisms with different genome sizes.

To assess the statistical significance of the calculated distances between the expression patterns of all genes within a trait, random background distributions are created for 1) individual genes and 2) traits of N genes. For individual genes, three different distributions were calculated, based on the distances between randomly sampled open reading frames, randomly sampled genes with an annotation (but not necessarily the same annotation), and randomly sampled genes with the same annotation. The background distribution for a trait of N genes is based on the distances between randomly composed sets of genes. For each gene pair, two distances metrics are calculated, the Pearson Correlation (PC) and the Normalized Rank Euclidean Distance (NRED). In practice, it is often found that a certain annotation is assigned to multiple genes in the same genome. If this occurs, there is an option to use either a random selection, or the highest scoring pair. In the latter case, a correction for multiple testing is implemented. This process is repeated N-times, where N equals the number of genes in any given trait. The background distribution for traits is determined by first randomly sampling two genomes, identifying the overlap in annotations, and finally artificially defining a trait containing N annotations. For each annotation in the trait, the distances are calculated between genome A and genome B, as described in the previous section. As modules vary in size, this process is repeated for traits of different sizes.

### Identifying Attributes

TbasCO provides both a cluster-based and pair-wise approach to identify attributes. In both methods, the distance between expression patterns of a trait between two genomes is first calculated based on a composite Z score of the PC and NRED for each gene composing the trait. In the cluster-based analysis, the distances are subsequently clustered using the Louvain clustering algorithm to identify trait attributes. To determine if an attribute is significantly similar or not, a one-sided T-test between the attribute and the random background distribution of traits is conducted. This is done for both cluster-based and model-based comparisons. Many traits are complex and represented in databases such as KEGG by numerous alternative routes. To deal with this complexity, each pathway is expanded into the Disjunctive Normative Form (DNF). Due to the extremely high number of DNFs for some traits, attributes are pruned based on a strict requirement of 100% completion.

### Distance Calculations

To determine the similarity in expression patterns between genes, two distance metrics are calculated: the PC between RNAseq counts across samples, and the NRED, where ranks are a measure of relative abundance of RNA in each sample, normalized the abundance of RNA in the corresponding genome. These distance scores are converted to Z scores using a background distribution of distances between randomly sampled genes as previously described. To determine statistically significant similarities between the expression patterns of a trait between two genomes, a composite distance score is calculated based on the distance between genes in two different genomes. For each of these genes the PC and NRED are calculated and transformed to Z scores and combined as (−1*PC + NRED). The distance of the trait between two genomes is defined as the average of these composite distance scores, and then normalized by the Jaccard distance between these genomes.

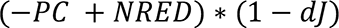

### Statistical Assessment of Trait Attributes

In both model-based and pair-wise approaches, the distance is first calculated between expression patterns of a trait between two genomes based on the composite Z score of the PC and NRED for each gene composing the trait. In the clustering-based analysis, the distances are subsequently clustered using the Louvain clustering algorithm to identify trait-attributes. To determine if attributes are significantly similar, a one-sided T-test is conducted between the attribute and a background distribution of randomly sampled traits with the same number of genes. To derive the random background distributions, multiple distributions are calculated ranging in gene numbers from the smallest trait to the largest trait in the dataset as described previously. For each background distribution, N (default: 10,000) traits are randomly composed. The distances between these artificial traits are calculated in the same way as for the actual traits. In addition to a statistical pruning step, the attributes are pruned based on a strict requirement of 100% completion of each DNF module. A benchmarking analysis to examine the effects of different parameters was conducted to determine their influence on the number of attributes identified and may be found in the supplementary materials (Supplementary Table 1, Supplementary Figures 2-4).

### Data and Code Availability

All supplementary files and figures including functional annotations and transcriptome count files are available at https://figshare.com/projects/EBPR_Trait-Based_Comparative_Omics/90437. All 64 flanking genomes have been deposited in NCBI at Bioproject <u>PRJNA714686</u>. The remaining two reassembled Accumulibacter genomes have not been deposited in NCBI to not confuse between the original CAPIA and CAPIIA assemblies [27, 28]. These contemporary assemblies are available at the Figshare repository. The three metagenomes and six metatranscriptomes used in this study are available on the JGI/IMG at accession codes 3300026302, 3300026286, 3300009517, and 3300002341-46, respectively. All code for performing metagenomic assembly, binning, and annotation can be found at https://github.com/elizabethmcd/EBPR-MAGs. The TbasCO method has been implemented as a reproducible R package and can be accessed at https://github.com/Jorisvansteenbrugge/TbasCO.

## RESULTS AND DISCUSSION

### Reconstructing a Diverse EBPR SBR Community

To explore trait-based transcriptional dynamics of a semi-complex microbial community, we applied genome-resolved metagenomics and metatranscriptomics to an EBPR sequencing-batch reactor (SBR) ecosystem (Figure 2). We previously performed a metatranscriptomics time-series experiment over the course of a normally operating EBPR cycle to investigate the regulatory controls of Accumulibacter gene expression [40]. In this experiment, six samples were collected for RNA sequencing: three from the anaerobic phase and three from the aerobic phase (Figure 2A). Additionally, three metagenomes were collected from the same month of the metatranscriptomic experiment, including a sample from the same date of the experiment. We reassembled contemporary Accumulibacter clade IIA and IA genomes that were previously assembled from the same bioreactor system [27, 28]. The genomes of Accumulibacter clades IA and IIA are similar by approximately 85% average-nucleotide identity [28], and although this is well below the common species-resolved cutoff of 95% [48], we refer to the clade nomenclature defined based on polyphosphate kinase (*ppk1*) sequence identity [49, 50]. During the experiment, the bioreactor was highly enriched in Accumulibacter clade IIA, accounting for approximately 50% of the mapped metagenomic reads and the highest transcriptional counts (Figures 2B and 2C) [40]. Whereas Accumulibacter clade IA exhibited low abundance patterns but was within the top 10 genomes with the highest total transcriptional counts (Figure 2C).

**Figure 2.**
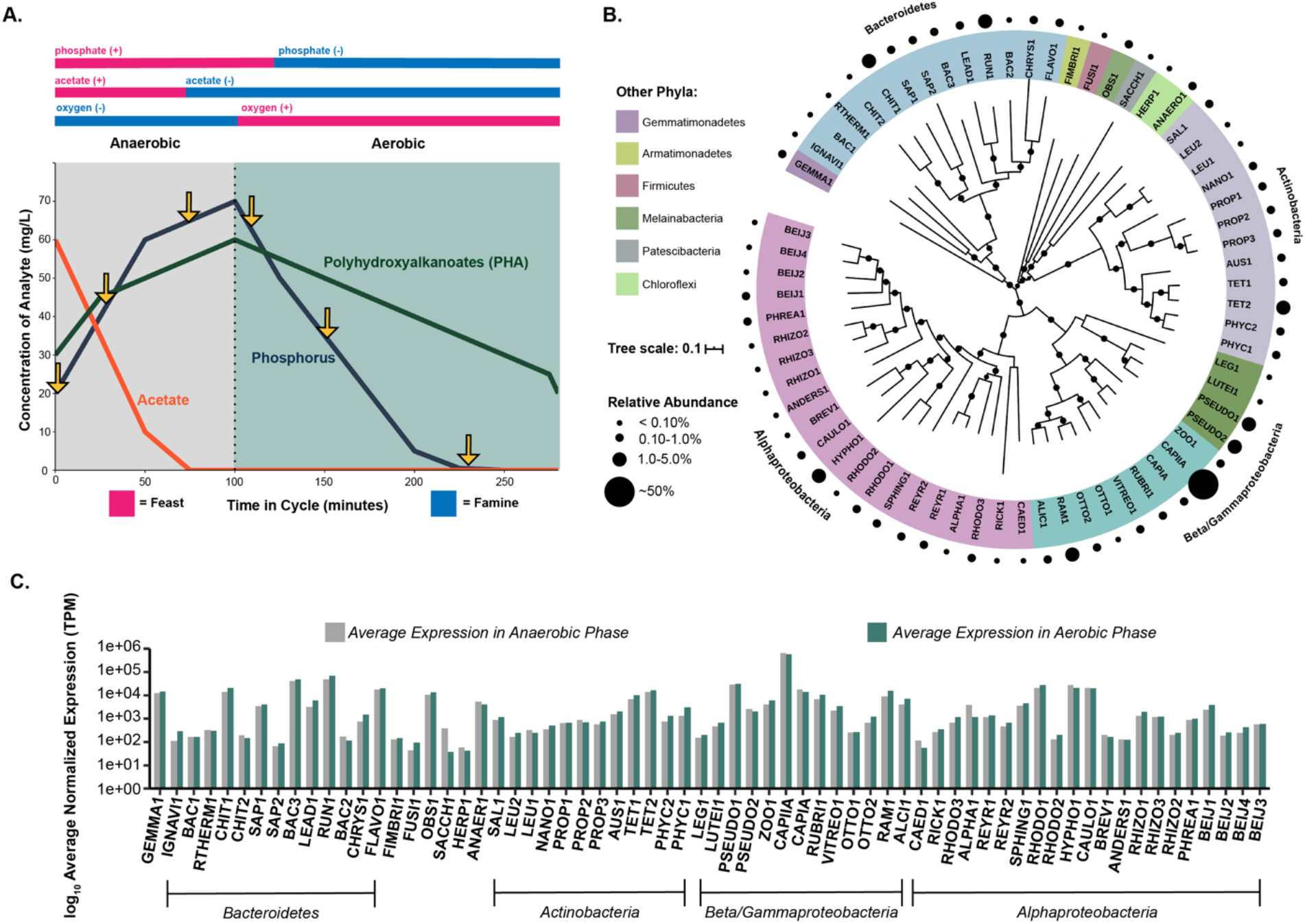
Genome-Resolved Metatranscriptomics Approach of an EBPR System. Application of a genome-resolved metatranscriptomics approach to a lab-scale sequencing batch reactor (SBR) designed to enrich for Accumulibacter. **1A)** Schematic of the main cycle parameters and analyte dynamics of an SBR simulating EBPR. Six samples were taken for RNA sequencing within the cycle at time-points denoted by arrows. **1B)** Phylogenetic identity and abundance patterns of 66 assembled MAGs from the EBPR system. The phylogenetic tree was constructed from concatenated markers contained in the GTDB-tk with muscle, calculated with RAxML, and visualized in iTOL. A phylogenetic tree of all 66 MAGs with reference genomes and high-quality genomes from Singleton et al. constructed with concatenated markers from GTDB-tk are provided in Supplementary Figure 1. Sizes of circles represent abundance patterns of metagenomic reads mapping back to genomes from the same day as the metatranscriptomic experiment and are not to scale. **1C)** Transcriptional patterns of each MAG in the anaerobic and aerobic phases of the EBPR cycle. RNA-seq reads from each time-point were competitively mapped to all 66 assembled MAGs and counts normalized by transcripts per million (TPM). Total counts in the anaerobic and aerobic phases for each genome were averaged separately and plotted on a log scale. Order of MAGs from left to right mirrors the order of MAGs in the phylogenetic tree in 1B from the top of the circle going clockwise.

Although this bioreactor system was highly enriched in Accumulibacter, a diverse flanking community persisted and was active in this ecosystem (Figure 2B, C). We reconstructed representative population genomes of the microbial community of the SBR system, resulting in 64 metagenome-assembled genomes (MAGs) of the flanking community. Interestingly, we recovered genomes of experimentally verified and putative PAOs, including two *Tetrasphaera spp.* (TET1 and TET2) ‘*Candidatus Obscuribacter phosphatis’* (OBS1), and *Gemmatimonadetes* (GEMMA1). Pure cultures of *Tetrasphaera* have been experimentally shown to cycle polyphosphate without incorporating PHA [36], deviating from the hallmark Accumulibacter PAO model. The first cultured representative of the *Gemmatimonadetes* phylum *Gemmatimonas aurantiaca* was isolated from an SBR simulating EBPR and was shown to accumulate polyphosphate through Neisser and DAPI staining [51]. Additionally, *Ca. Obscuribacter phosphatis* has been hypothesized to cycle phosphorus based on the presence of genes for phosphorus transport, polyphosphate incorporation, and potential for both anaerobic and aerobic respiration [37], and has also been enriched in photobioreactor EBPR systems [52]. Both *Tetrasphaera spp.* TET1 and TET2, OBS1, and GEMMA1 groups exhibit higher relative abundance patterns than CAPIA but have similar relative transcriptional levels (Figure 2B and 2C, Table 1).

Numerous SBR MAGs among the *Actinobacteria* and *Proteobacteria* contain the metabolic potential for phosphorus cycling based on the presence of the high-affinity phosphorus transporter *pstABCS* system, polyphosphate kinase *ppk1*, and the low-affinity *pit* phosphorus transporter (Supplementary Figure 5). Additionally, select MAGs within the *Alphaproteobacteria*, *Betaproteobacteria*, and *Gammaproteobacteria* contain all required subunits for polyhydroxyalkanoate synthesis (Supplementary Figure 5). Other abundant and transcriptionally active groups in the SBR ecosystem that are not predicted to be PAOs are members of the *Bacteroidetes* such as CHIT1 within the *Chitinophagaceae,* and *Cytophagales* members *Runella* sp. RUN1 and *Leadbetterella* sp. LEAD1 (Figure 2B and 2C, Table 1). Interestingly, an uncharacterized group within the *Bacteroidetes* BAC1 contributed the third most to the pool of transcripts (Figure 2C), and did not show phylogenetic similarity to MAGs assembled from Danish full-scale wastewater treatment systems [39] (Supplementary Figure 1). Other groups from which we assembled MAGs for that do not exhibit clear roles in EBPR systems were *Chloroflexi* ANAER1 and HERP1 MAGs, *Armatimonadetes* FIMBRI1, *Firmicutes* FUSI1, and *Patescibacteria* SACCH1. Members of the *Chloroflexi* are filamentous bacteria that have been associated with bulking and foaming events in full-scale WWTPS [53–55], but also aid in forming the scaffolding around floc aggregates and degrade complex polymers [55–57]. The *Patescibacteria* (formerly TM7) are widespread but low abundant members of natural and engineered ecosystems, contain reduced genome sizes, and may contribute to filamentous bulking in activated sludge [21, 58]. To summarize, lab-scale SBRs designed to enrich for Accumulibacter contain diverse flanking community members [27, 32], but their ecological functions and putative interactions remain to be fully understood in the context of the EBPR ecosystem.

### Identifying Expression-Based Trait Attributes Among the EBPR SBR Community with TbasCO

Current metatranscriptomics approaches often employ either a gene-centric [31, 59–61] or genome-centric approaches [40, 62–64]. In both approaches, highly, differentially, or co-expressed genes are identified and tested for enrichment of specific functions. Enrichment- or annotation-based approaches are employed in numerous metatranscriptomics tools such as MG-RAST, MetaTrans, SAMSA2, COMAN, IMP, and Anvi’o [65–70]. Here, we expand on the use of molecular markers as traits by defining expression attributes by leveraging *a priori* knowledge from predefined trait libraries, such as the KEGG database [71], to statistically assess inter-species expression patterns of genes that together form a trait (Figure 1). First, our results showed that there is statistically significant transcriptional conservation of genes at the community level; genes that share an annotation were significantly more similar than expected using two different distance metrics (NRED: p-value < 2.2e-16, PC: p-value < 2.2e-16). Extending this statistical analysis to the trait level, we identified 1674 attributes distributed across the 66 genomes. On average, we identified 9.12 genomes per attribute (SD - 5.22), with a minimum of 3 genomes and a maximum of 35 (Figure 3A). Based on these statistics, we defined redundant attributes as those two standard deviations above the mean (19 genomes). With this cutoff applied, we identified 79 redundant trait attributes mostly belonging to pathways among carbohydrate metabolism, purine metabolism, and fatty acid metabolism categories (Table 2). Of 290 traits, we identified 97 traits with two or more attributes identified (33%). Of these, traits in 10 or more genomes were twice as likely to have two or more attributes (65%), suggesting that divergent expression patterns for a trait are common, and may represent a niche-differentiating feature (Figure 3A). Henceforth, when multiple attributes are identified for a trait, we refer to these as niche-differentiating attributes.

**Figure 3.**
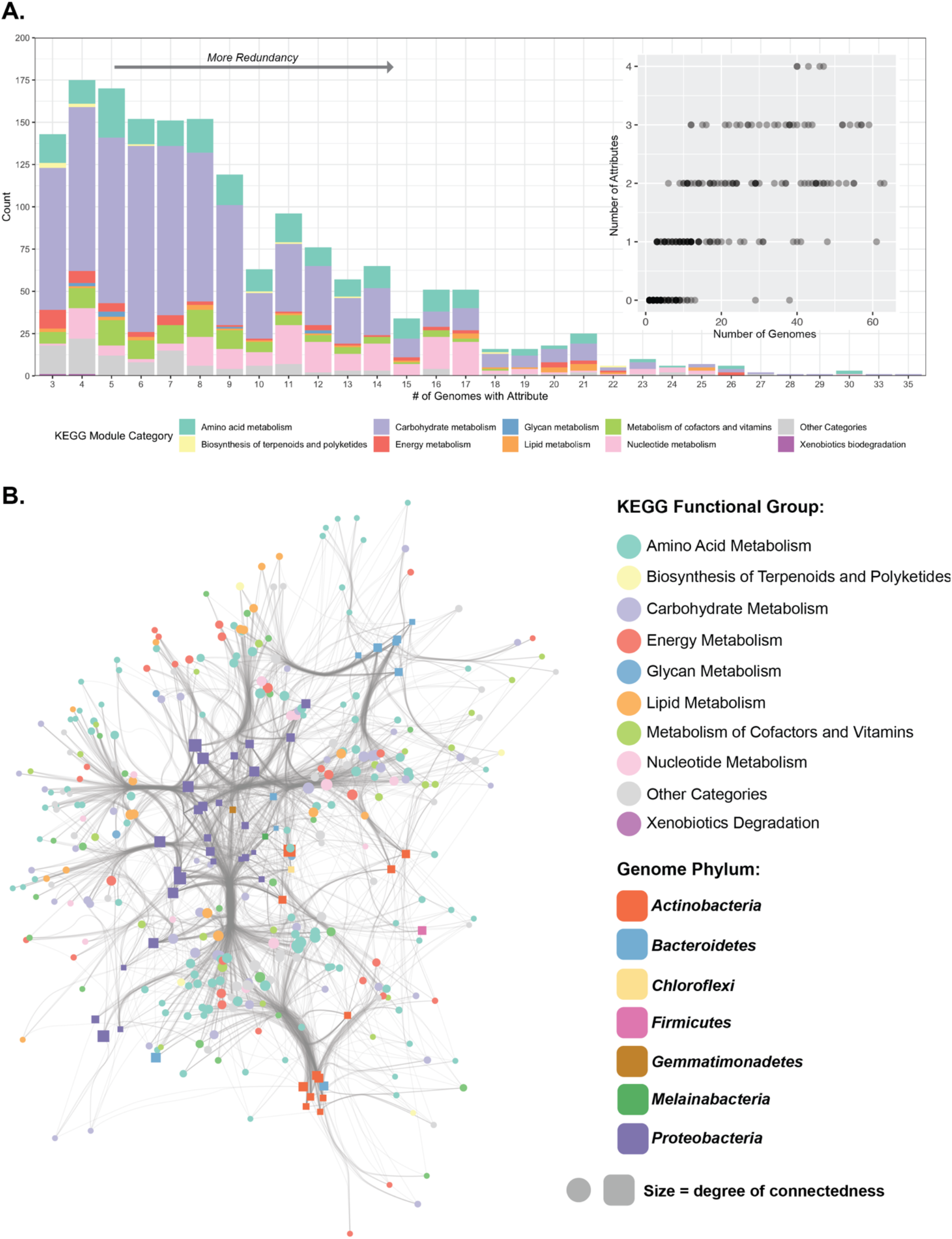
Clustering and Distribution of Trait Attributes Across EBPR SBR Community Members. Using the TbasCO method, we identified expression-based trait attributes from predefined trait modules in the KEGG library and explored the distribution of these trait attributes across community members. **A)** Distribution of trait-attributes among sets of genomes. Bars represent the number of trait-attributes present in a set number of genomes and colored by KEGG module category. Among a total of 35 genomes, trait attributes present between 3-18 genomes are designated as niche differentiating, whereas trait attributes present in 19 or greater genomes are designated as core trait attributes. Inset figure demonstrates the maximum number of attributes for the maximum number of genomes. **B)** Cytoscape network showing the connectedness of genomes to trait attributes. The network was filtered to only include nodes with more than 5 connections, therefore filtering out both genomes with few trait attributes and trait attributes connected to less than 5 genomes. Genomes are represented as squares colored by phylum, and trait attributes are represented as circles colored by KEGG category. The size of both the squares and circles represents the number of connections to that genome or trait attribute, respectively.

**Table 2.**
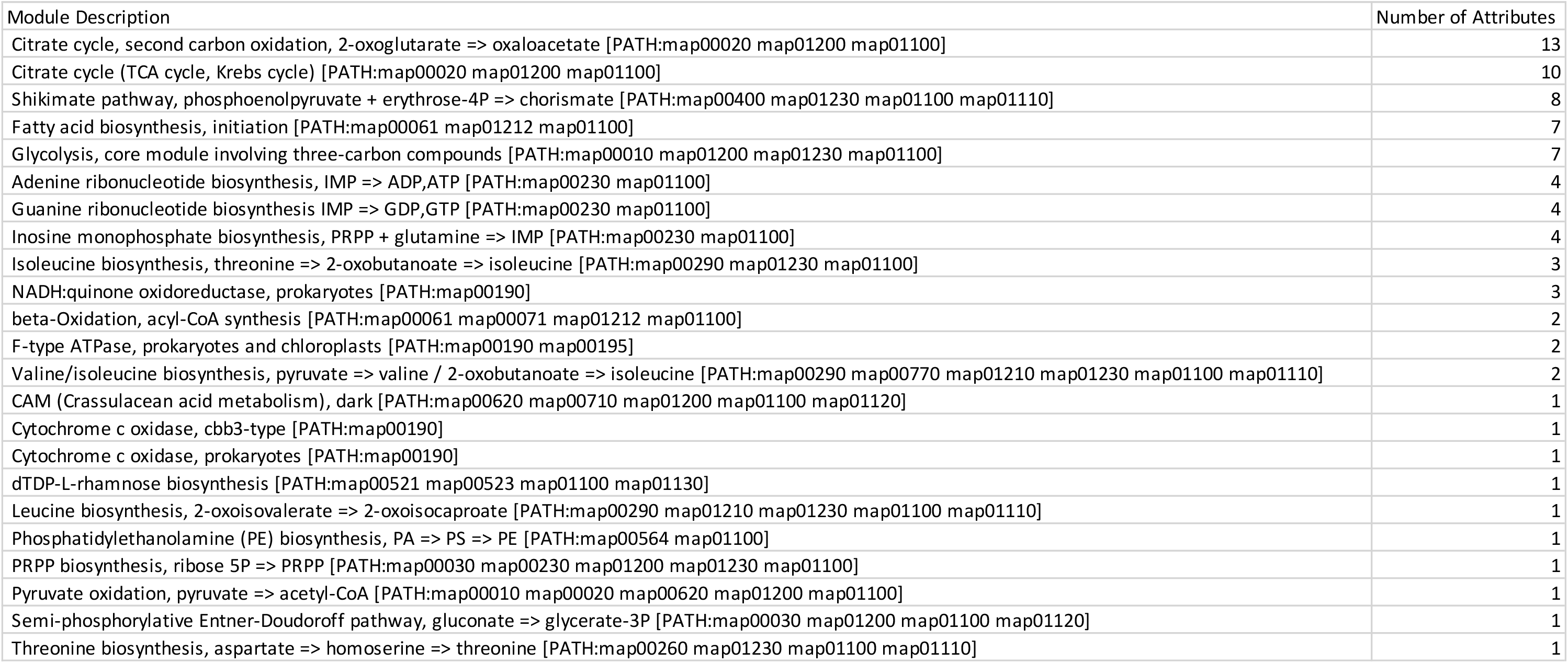
KEGG Pathways for core trait-attributes present in greater than 19 genomes.

From the ecosystem perspective, a clear phylogenetic signal is observed in the distribution of attributes, as genomes cluster together by shared trait attributes by phylum with some exceptions, such as genomes belonging to the *Bacteroidetes, Actinobacteria,* and *Proteobacteria* clustering together, respectively (Figure 3B). For simplicity, we filtered the network to only include nodes with more than 5 connections. Highly redundant trait attributes belonged to modules in the lipid metabolism, energy metabolism, and nucleotide metabolism KEGG functional categories. In contrast, more specialized trait attributes on the periphery of the network or amongst group-specific clusters such as within the *Actinobacteria* or subsets of the *Proteobacteria* belonged to amino acid metabolism, biosynthesis of terpenoids and polyketides, metabolism of cofactors and vitamins, and carbohydrate metabolism KEGG modules. Pathways of note that showed a high level of redundancy include the TCA cycle, isoleucine biosynthesis, acyl-CoA synthesis, threonine biosynthesis, and cytochrome c oxidase activity (Table 2). Large pathways with hundreds of possible routes such as glycolysis, the TCA cycle, gluconeogenesis, and the pentose phosphate pathway are not included in the main network and are displayed as individual networks (Supplementary Figure 6).

We next explored the distribution of non-redundant attributes (e.g. 3-18 genomes) (Figure 3A). A total of 796 trait attributes with low redundancy were identified belonging to pathways involved in carbohydrate cofactor and vitamin metabolism including glycolysis, gluconeogenesis, parts of the TCA cycle, tetrahydrofolate biosynthesis, tryptophan biosynthesis, and the pentose phosphate pathway (Table 3). Different sets of low redundancy trait attributes were identified within respective phyla (Supplementary Figure 7). Between genomes belonging to the *Actinobacteria*, *Alphaproteobacteria, Bacteroidetes, Betaproteobacteria,* and *Gammaproteobacteria,* low redundancy attributes (belonging to less than half of the total genomes within the phylum) include carbohydrate metabolism, amino acid metabolism and metabolism of cofactors and vitamins (Supplementary Figure 7). Redundant trait attributes within individual phyla belong to core energy metabolism pathways, fatty acid biosynthesis, and carbohydrate metabolism. However, even within individual phyla, non-redundant attributes include different amino acids and cofactors (Extended Table 1 - available on Figshare https://figshare.com/articles/dataset/Lineage-Specific_Core_and_Niche_Differentiating_Traits/15001200).

**Table 3.**
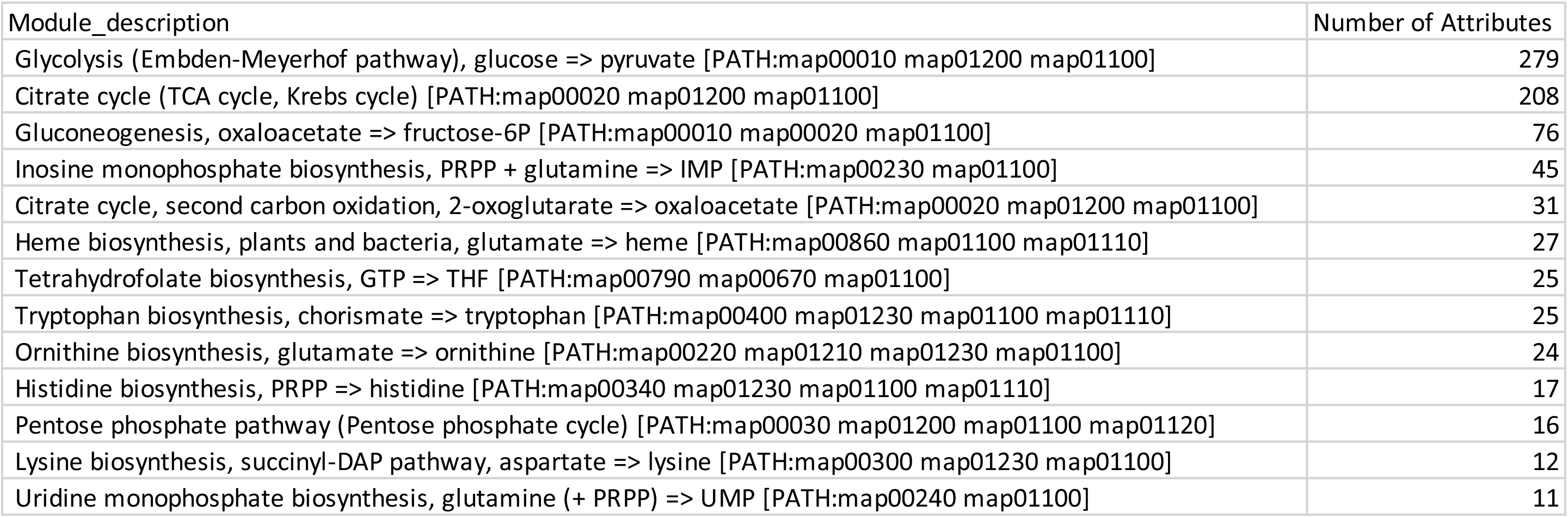
KEGG Pathways for differentiating trait-attributes present between 3 and 18 genomes.

As noted previously, one of the most striking findings is that a majority, 65% of traits present in 10 or more genomes have multiple expression attributes. Thus, it seems that while the presence of marker genes suggests many organisms share a particular trait, the presence of niche-differentiating expression profiles suggest an alternative story, that there is a level of hidden metabolic diversity. For example, central carbon metabolism and energy pathways such as the TCA cycle, glycolysis, gluconeogenesis, and the pentose phosphate pathway are oftentimes considered core traits when only analyzing the presence and/or absence of individual markers belonging to these pathways. Among over 1000 high-quality MAGs assembled from full-scale Danish WWTPs, the TCA cycle and pentose phosphate pathway are highly represented among the abundant microorganisms, with glycolysis less so [39]. Whereas the TCA cycle and pentose phosphate pathway are present among a high number of genomes in the EBPR SBR community, different routes or parts of these pathways have niche-differentiating distributions (Supplementary Figure 4, Tables 2 and 3). These finer-scale differences in expression of “core” traits may explain the persistence of a diverse community when solely fed acetate, as different lineages could employ similar carbon utilization pathways differently or in more versatile ways. Another salient aspect of this analysis is the astonishingly high number of possible routes within individual pathways here represented by their Disjunctive Normal Forms. For example, accounting for all alternative routes and enzymes, the glycolysis pathway has 100s of possible routes. Layering upon this many expression attributes reveals a large hidden metabolic versatility.

### Dimensionality of the High-Affinity Phosphorus Transporter System *PstABCS*

The EBPR ecosystem is characterized by its highly dynamic phosphorus cycles. To explore how different lineages respond to fluctuating phosphorus concentrations, we explored the expression-based attributes for the KEGG module of the high-affinity phosphorus transporter *pstABCS* (Figure 4). The *pstABCS* system is an ABC-type transporter that strongly binds phosphate under phosphorus-limiting conditions; therefore, it would be expected that the highest expression levels would be at the end of the aerobic cycle [72]. In contrast, we found that expression of the *pstABCS* was characterized by two different trait attributes. In the first attribute shared by 14 community members, all components of *pstABCS* displayed the highest activity towards the end of the aerobic cycle, when phosphorus concentrations were depleted (Figure 4, Attribute 1). Conversely, 11 community members displayed an alternate attribute where the highest activity of *pstABCS* was at the transition from anaerobic to aerobic phases when phosphorus concentrations are highest (Figure 4, Attribute 2).

**Figure 4.**
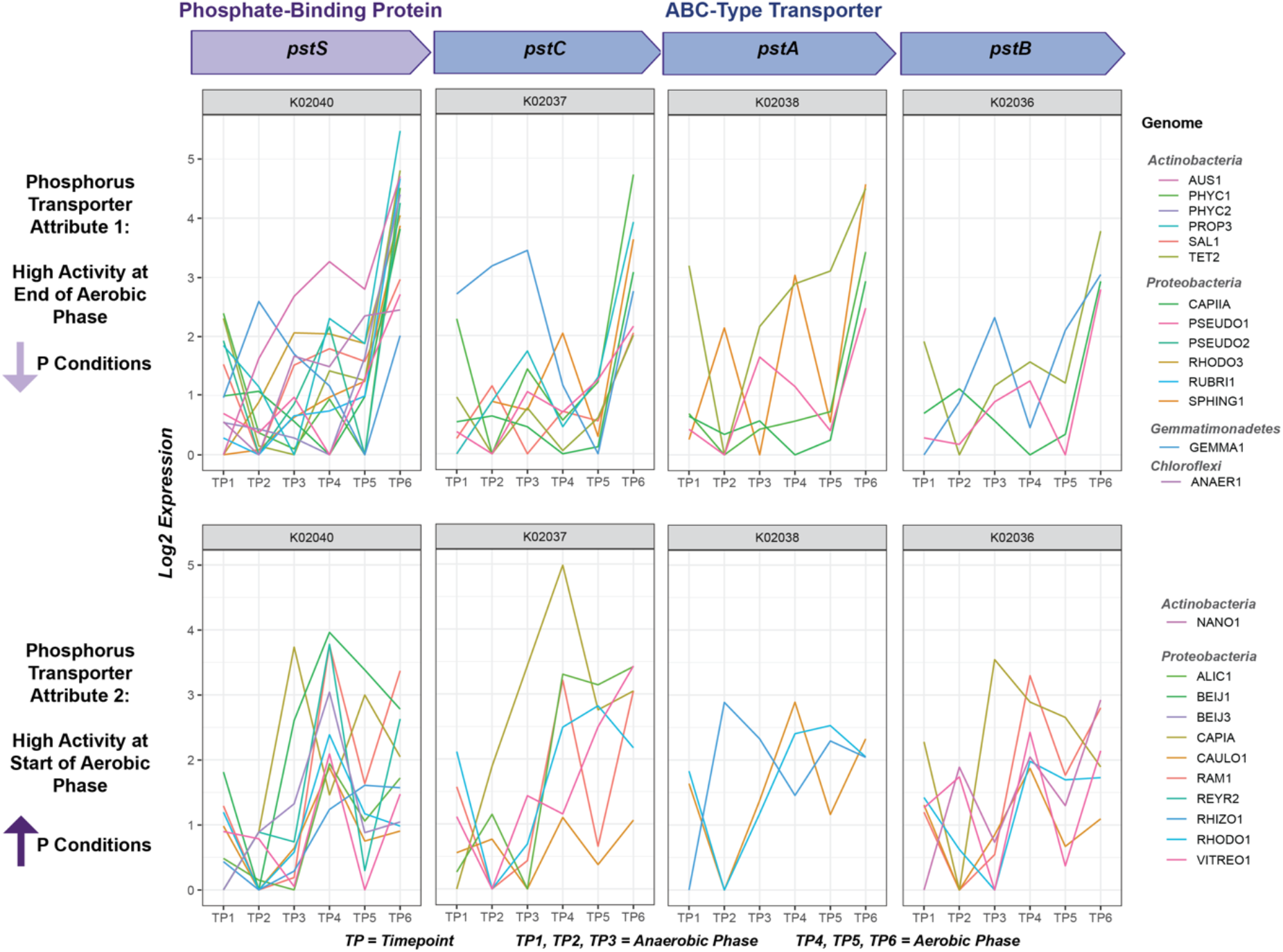
Trait Attributes of the High-Affinity Phosphorus Transporter System *pstABCS*. Using the TbasCO method, two trait attributes of the high-affinity phosphorus transporter system *pstABCS* were identified. The *pstABCS* system consists of a phosphate-binding protein and ABC-type transporter, and the corresponding KEGG orthologs for each subunit are shown. Timepoints 1-3 refer to the three anaerobic phase timepoints, and timepoints 4-6 refer to the three anaerobic phase timepoints (Figure 1). Expression values are log-transformed based on setting the lowest expression value within each genome across the time-series to 0 for each subunit. Specific subunits for some genomes in both attributes are missing to the high cutoff thresholds for annotations. However we kept genomes with 2/4 subunits to show similarities in expression profiles. The first *pstABCS* trait-attribute includes microbial lineages that exhibited the highest expression of all subunits towards the end of the aerobic cycle, when phosphate concentrations are expected to be lowest. This includes microbial lineages within the *Actinobacteria, Proteobacteria, Gemmatimonadetes,* and *Chloroflexi.* The second *pstABCS* trait-attribute includes lineages that exhibited highest expression of all subunits upon the switch from anaerobic to aerobic phases, or when phosphate concentrations are expected to be the highest. This includes lineages within the *Actinobacteria* and *Proteobacteria*.

These results are in agreement with previous results showing that Accumulibacter clade IIC has a canonical *pstABCS* expression pattern (as in Figure 4, Attribute 1) , whereas the Accumulibacter clade IA has a non-canonical expression (as in Figure 4, Attribute 2) [31]. By assigning trait attributes, we are able to extend these findings beyond Accumulibacter to other flanking community members in the SBR ecosystem suggesting that there are conserved ecological pressures driving niche differentiating expression patterns in *pstABCS* within the EBPR community.

### Distribution and Expression of Truncated Denitrification Steps Among EPBR Community Members

Understanding the induction of denitrification is an important ecosystem property linked to the redox status of an environment. In EBPR communities, there are many diverse and incomplete denitrification pathways, distributed across many lineages denitrification steps expected in denitrifying systems (Figure 5). Among all 66 MAGs, we did not identify any single MAG with a complete denitrification pathway consisting of the genetic repertoire necessary to fully reduce nitrate to nitrogen gas (Supplementary Figure 5). Instead, we identified multiple groups of organisms with truncated denitrification pathways, with steps distributed among cohorts of community members (Figure 5).

**Figure 5.**
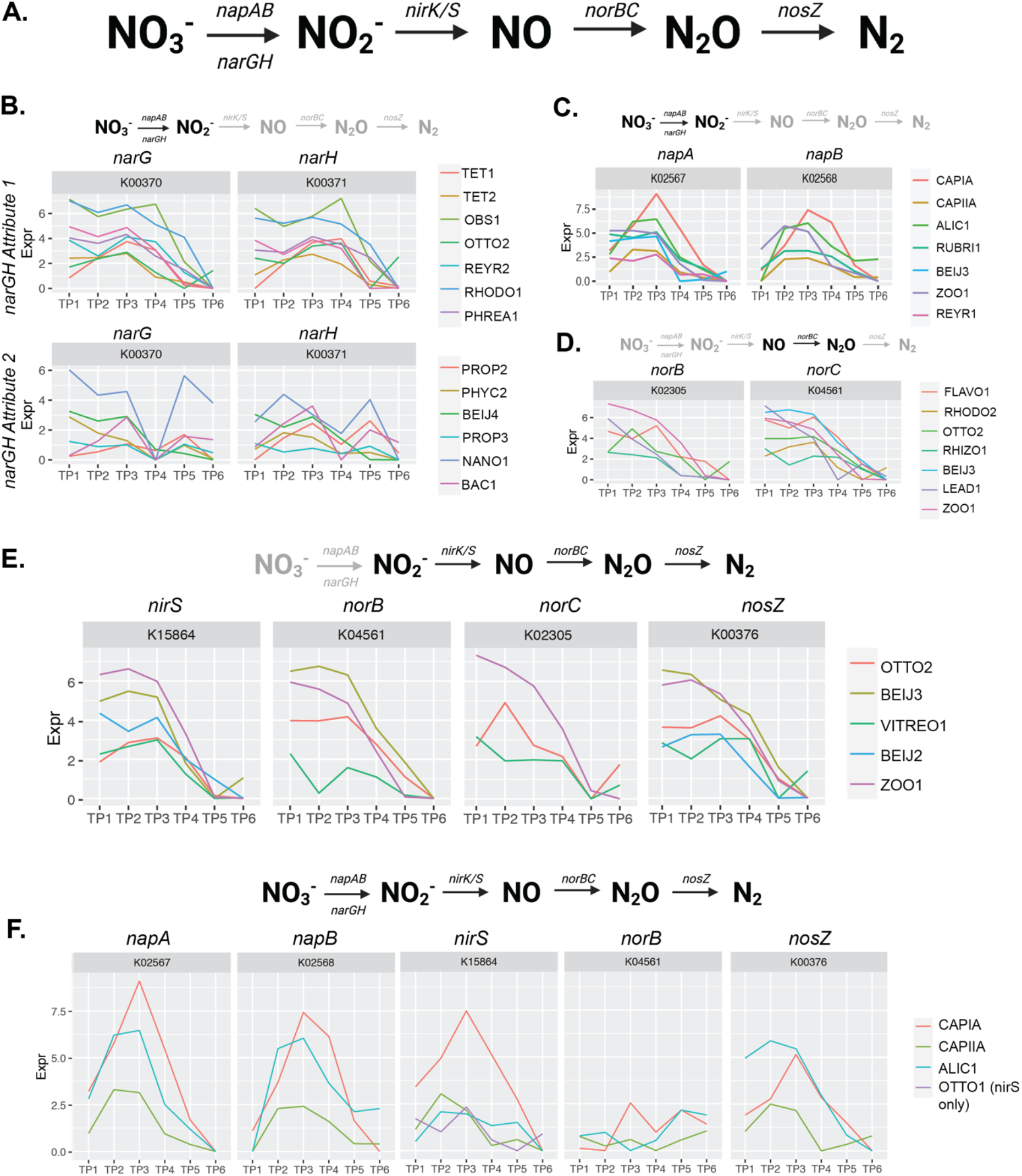
Expression Dynamics of Distributed Denitrification Routes. Expression of denitrification traits distributed among community members in the EBPR SBR ecosystem. Timepoints 1-3 correspond to the anaerobic phase and timepoints 4-6 correspond to the aerobic phase as referenced in Figure 1. **A)** Complete denitrification pathway and associated genetic repertoire with each sequential step. **B)** Trait attributes of expression dynamics for community members with the *narGH* nitrate reductase system. This trait was the only denitrification trait identified with more than one attribute. **C)** Expression dynamics of the *napAB* nitrate reductase system. **D)** Expression dynamics of the *norBC* nitrous oxide reductase system. **E)** Expression of all steps of denitrification starting at nitrite reduction. **F)** Expression of the most complete denitrification route among three community members, with the *norC* subunit for nitrous oxide reduction missing. Note that OTTO1 only contains *nirS* but is included in this trait attribute because the expression dynamics are similar to that of the other three genomes for this subunit.

For the first steps of reducing nitrate to nitrite, we explored expression attributes of the *napAB* and *narGH* pathways (Figure 5B, C). For the *narGH* pathway, two attributes were identified (Figure 5B). The first *narGH* attribute was characterized by high expression in the anaerobic phase, with decreasing activity by the second time point of the anaerobic phase. Genomes containing this attribute included the experimentally verified and putative PAOs *Tetrasphaera* (TET1 and TET2) and *Ca.* Obscuribacter (OBS1), respectively. The second attribute was exhibited among members of the *Actinobacteria* (PROP2, PHYC2, PROP3, and NANO1), *Proteobacteria* (BEIJ4), and *Bacteroidetes* (BAC1). The attribute identified for *napAB* was also more highly expressed anaerobically and included CAPIA, CAPIIA, ALIC1, REYR2, RUBRI1, and BEIJ3. Interestingly, this *napAB* attribute had expression patterns that quickly decreased in the first aerobic time point, suggesting a tighter regulation than Attribute 1 for *narGH*. Together, this suggests that the regulation of denitrification within the EBPR ecosystem is a niche-differentiating feature whereby the induction of denitrification pathways occurs either anaerobically or only after anaerobic carbon contact.

A smaller cohort contained the genetic repertoire to reduce nitrite to nitrogen gas and exhibited hallmark anaerobic-aerobic expression patterns (Figure 5E) These members within the *Proteobacteria* (OTTO2, BEIJ3, VITREO1, and ZOO1) contained the *nirS* nitrite reductase, the *norBC* nitric oxide reductase, and *nosZ,* and showed highest expression of these subunits towards the beginning of the anaerobic cycle, slowly decreasing over the aerobic period to their lowest in the end of the aerobic cycle. Although BEIJ2 was lacking the *norBC* system, it contained the *nirS* nitrite reductase and *nosZ* subunit, and exhibited similar expression patterns to others in this cohort. Other *Proteobacteria* lineages only contained the *norBC* subunits but were expressed in similar fashions (RHODO2, FLAVO1, RHIZO1, and LEAD1) (Figure 5D). Accumulibacter clades IA and IIA as well as ALIC1 were the only lineages with near-complete denitrification pathways. These lineages contained the *napAB* nitrate reductase system as mentioned above, the *nirS* nitrite reductase, *norB* (missing a confident hit for the *norC* subunit), and *nosZ.* These three lineages also exhibited hallmark upregulation of all steps in the anaerobic phase, with decreased activity after aerobic contact (Figure 5F).

Interestingly, Accumulibacter clade IA exhibited a higher magnitude of expression of denitrification steps when activity levels were normalized relative to clade IIA, supporting the hypothesis that denitrification is a niche-differentiating feature among clades [28, 31, 73], and possibly a strain-specific trait since denitrification traits cannot be predicted based on *ppk1* clade designations [32]. For example, independent observations in differences among denitrification activities among strains within Accumulibacter clade IC are inconsistent [34, 74]. Within the same bioreactor environment, coexisting Accumulibacter clades differ between denitrification abilities and expression profiles [31, 33, 75]. Truncated denitrification pathways have also been previously shown to be distributed among community members, with the complete denitrification genetic repertoire only present in few members [33, 75], which could be due to extensive horizontal gene transfer of genes comprising denitrification steps [75, 76]. Although this experiment was not conducted under denitrifying conditions, our approach could be applied to denitrifying EBPR systems to further understand the distribution of denitrification traits among community members and how to selectively enrich for diverse DPAOs.

### Biosynthetic Potential and Expression Dynamics of Amino Acid and Vitamin Synthesis Pathways

Although SBRs are designed to enrich for Accumulibacter by providing acetate as the sole carbon source, a diverse flanking community persists in these setups [27, 75]. One hypothesis for the persistence of flanking community members may be cooperative interactions due to underlying auxotrophies of amino acid and vitamin biosynthetic pathways in Accumulibacter. Amino acids and vitamin cofactors are metabolically expensive to synthesize, and widespread auxotrophies have been widely documented among microbial communities [77, 78]. Specifically, auxotrophies of vitamin cofactors have been shown to fuel bacterial and cross-kingdom interactions with *de novo* bacterial and cross-kingdom interactions with *de novo* synthesizers [79, 80]. To explore this hypothesis in the EPBR SBR community, we analyzed the presence of amino acid and vitamin biosynthetic pathways and their expression patterns among the top 15 genomes based on transcript abundance (Figure 6).

**Figure 6.**
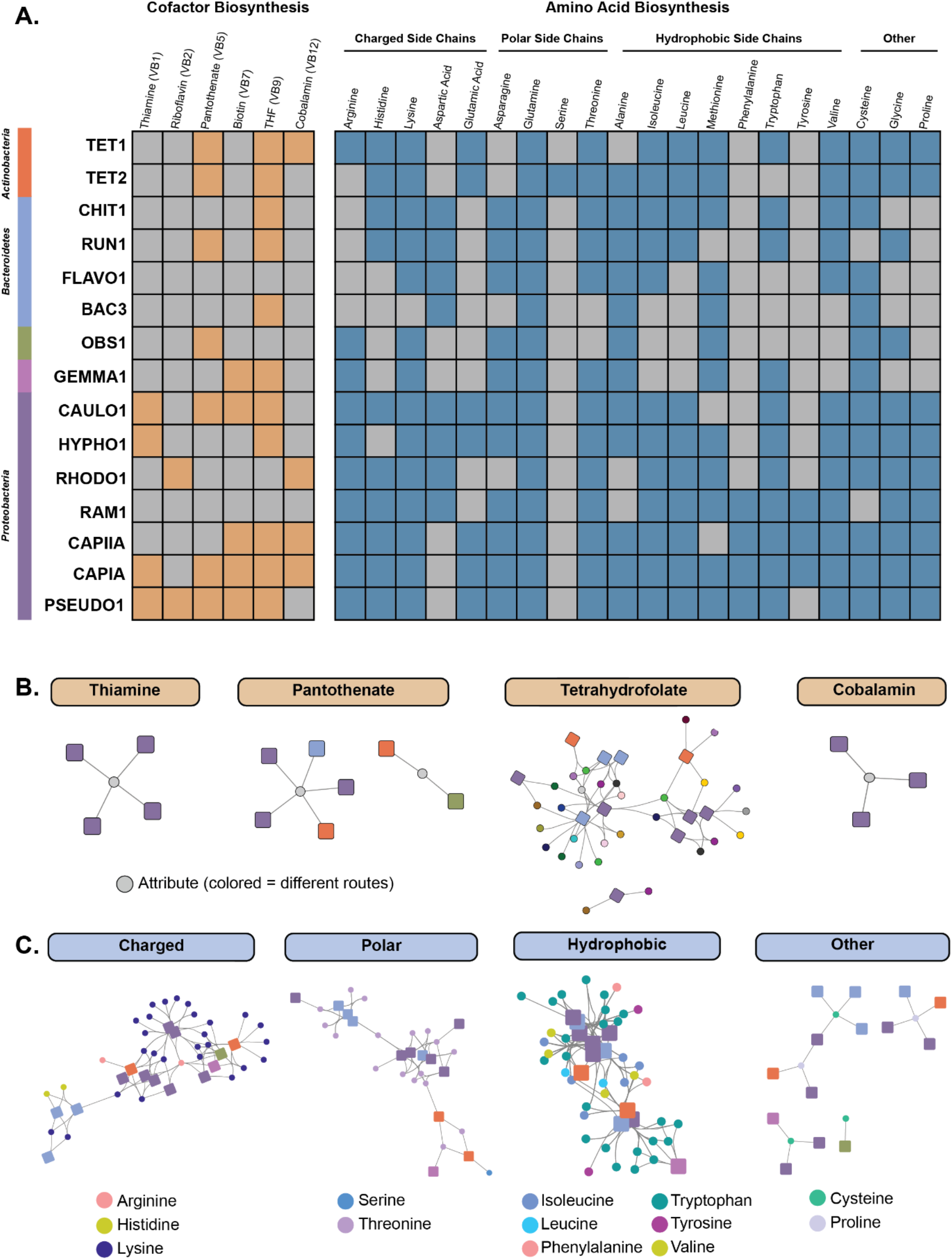
Biosynthetic Potential Compared to Expression of Amino Acid and Vitamin Synthesis Pathways for Top 15 Expressed MAGs. Biosynthetic potential and expression patterns of amino acid and vitamin pathways were analyzed for the top 15 genomes with the highest transcriptional counts (Table 1). **A)** For a pathway to be considered present for downstream analysis in the TbasCO pipeline, 80% of the pathway had to be present in a genome. Thus, we used this cutoff criterion to discern whether a specific pathway was present or absent in a genome (with the expectation of methionine, as all genomes did not contain at least 80% of the subunits in the KEGG methionine synthase pathway, we inferred the presence of the methionine synthase as presence of this pathway). Orange colored boxes for cofactor biosynthesis pathways represents the presence of that pathway, whereas grey infers absence. For amino acid biosynthetic pathways, amino acids are listed by their side chain groups – charged, polar, hydrophobic, and other. **B)** Mini-networks of vitamin co-factors. Squares are genomes with the colors matching the color bar in A. Nodes are attributes, where the colored nodes for the tetrahydrofolate attributes represent the different routes. **C)** Mini-networks of amino acid biosynthesis pathways split by type. Colors of nodes for each amino acid represent the different routes for that pathway. Squares represent genomes with colors matching the color bar in A.

Within Accumulibacter, there are a few key vitamin cofactor and amino acid auxotrophies that could fuel potential interactions with flanking community members. Both Accumulibacter clade genomes are missing the riboflavin pathway for FAD cofactor synthesis, as well as the pathways for serine and aspartic acid (Figure 6A). The biosynthetic pathway for aspartic acid is distributed among members of the *Bacteroidetes* and *Proteobacteria*, whereas only TET2 contains the pathway for serine synthesis (Figure 5A). The lack of serine biosynthesis pathways in Accumulibacter and other flanking genomes seems striking given that serine is one of the least metabolically costly amino acids to synthesize [81]. Interestingly, Accumulibacter clade IIA does not contain the biosynthetic machinery for thiamine and pantothenate synthesis, whereas clade IA does (Figure 6A). Only the CAULO1, HYPHO1, and PSEUDO1 genomes within the *Proteobacteria* can synthesize thiamine, whereas several other members can synthesize pantothenate (Figure 6A). The absence of the pantothenate biosynthetic pathway in Accumulibacter CAP IIA is particularly interesting given that coenzyme A is essential for polyhydroxyalkanoate biosynthesis, which fuels the polymer cycling PAO phenotype of Accumulibacter [24].

In addition to flanking community members potentially supporting the growth of Accumulibacter due to underlying auxotrophies, the reciprocal logic may be possible as well. Both Accumulibacter clades contain the pathways for synthesizing tyrosine and phenylalanine, which are missing in a majority of the top 15 active flanking genomes (Figure 6A). Only two other members within the *Proteobacteria* can synthesize tyrosine and phenylalanine, where RAM1 can synthesize both and PSEUDO1 only phenylalanine. Interestingly, phenylalanine and tyrosine are the second and third most metabolically expensive amino acids to synthesize, respectively, with tryptophan the most costly [81]. Additionally, a few highly active flanking community members lack the biosynthetic machinery for several vitamin cofactors and amino acids, such as FLAVO1 and BAC3 within the *Bacteroidetes* and the putative PAO *Ca.* Obscuribacter phosphatis OBS1 (Figure 6A). Particularly, RAM1 within the *Proteobacteria* is missing the biosynthetic machinery for all vitamin cofactors but can synthesize most amino acids including the most metabolically expensive as mentioned above.

We next analyzed the distribution of trait-attributes of vitamin and amino acid pathways among these genomes to understand how these biosynthetic pathways are expressed similarly or differently in the EBPR SBR ecosystem (Figure 6B and C). Members of the *Proteobacteria* containing thiamine and cobalamin biosynthetic pathways all express these traits similarly (Figure 6B). However, the pantothenate synthesis pathway contains two trait-attributes and is expressed differently among two cohorts. In the first attribute, RUN1, TET1, CAULO1, CAPIA, and PSEUDO1 express the pantothenate pathway similarly. However, OBS1 and TET2 express the pantothenate pathway differently (Figure 6B). Because tetrahydrofolate can be synthesized through different metabolic routes, we analyzed the differences in trait attribute expression for all routes in genomes that contained sufficient coverage of this trait. Members of the *Bacteroidetes* and *Proteobacteria* mostly cluster together among tetrahydrofolate attributes, whereas the TET1 and TET2 genomes are differentiated (Figure 6B).

Expression of various groups of amino acids show more differentiated patterns of expression for genomes with these pathways. Several amino acids also contain different metabolic routes for biosynthesis, and we analyzed all trait attributes for each amino acid for all routes grouped by type (Figure 6C). For the charged amino acids arginine, histidine, and lysine, members of the *Proteobacteria* and *Bacteroidetes* cluster within their phylogenetic groups, respectively, with lysine and histidine expressed differently among these groups (Figure 6C). In contrast, arginine is expressed similarly among all *Proteobacteria* genomes. Among the polar charged amino acids, TET2 is the only genome among the top 15 genomes that contains the metabolic pathway to synthesize serine (Figure 6A). Several groups contain the pathway for threonine synthesis, and expression of different threonine routes are differentiated among the *Proteobacteria, Bacteroidetes,* and *Tetrasphaera spp.,* but mostly clusters phylogenetically (Figure 6C). Notably, the expression patterns for the cysteine and proline biosynthetic pathways do not cluster phylogenetically, such as both *Tetrasphaera* genomes expressing the proline pathway more similarly to other *Proteobacteria* and *Bacteroidetes* (Figure 6C). The few lineages that can synthesize tyrosine and phenylalanine (CAPIA, CAPIIA, RAM1, PSEUDO1) show different patterns of expression. These results show that beyond the presence or absence of key vitamin cofactor and amino acid biosynthetic pathways, EBPR SBR organisms also display coherent and differentiated patterns of expression for these traits, of which the functional consequences remain to be further understood.

## CONCLUSIONS AND FUTURE PERSPECTIVES

In this work, we applied a novel trait-based ‘omics pipeline to a semi-complex, engineered bioreactor microbial community to explore ecosystem-level and niche-differentiating traits. Through assembling high-quality MAGs of the EBPR SBR community and using a time-series metatranscriptomics experiment, we were able to extend functional predictions and ecosystem inferences beyond hypotheses made from gene presence/absence data. Using our novel trait-based comparative ‘omics pipeline, we identified how similarities and differences in the expression of significant EBPR traits are conferred among community members such as phosphorus cycling, denitrification, and amino acid metabolism. Specifically, we demonstrate that traits with similar expression profiles may be clustered into attributes providing a new layer to trait-based approaches.

We believe that identifying expression-based attributes will be a powerful tool to explore microbial traits in natural, engineered, and host-associated microbiomes. Outside of activated sludge systems, trait-based approaches could illuminate how similar secondary metabolite clusters are expressed among different species in a community [82, 83], how auxotrophies for amino acid and vitamin cofactors govern interactions [84], how rhizosphere microorganisms respond to day-night cycles, and identify putative traits that universally exhibit ecosystem-level or niche-differentiating patterns across ecosystems [19, 23]. Importantly, our trait-based approach can be used to screen for expected expression patterns of a key trait compared to a model organism, and then prioritize specific microbial lineages for downstream experimental verification with techniques such as Raman-FISH [85, 86]. Overall, our trait-based comparative ‘omics pipeline is a novel and high-throughput approach to understand how microbial traits connect to ecosystem-level processes in diverse microbiomes.

## ACKNOWLEDGEMENTS

We thank Caitlin Singleton for providing early access to high-quality genomes from a full-scale WWTP to compare our MAGs against. Metagenomic and metatranscriptomic sequencing was provided through a Joint Genome Institute Community Science Proposal (Proposal ID 873). This work was supported by funding from the National Science Foundation (MCB-1518130) to K.D.M and D.R.N. Funding was provided to E.A.M. by a fellowship through the Department of Bacteriology at the University of Wisconsin – Madison. Funding for B.O.O was in part provided by the Technology Foundation of the Dutch National Science Foundation (NWO-TTW). This research was performed in part using the Wisconsin Energy Institute computing cluster, which is supported by the Great Lakes Bioenergy Research Center as a part of the U.S. Department of Energy Office of Science (DE-SC0018409).

