## Supplementary Materials for "TbasCO: Trait-based Comparative ’Omics Identifies Ecosystem-Level and Niche-Differentiating Adaptations of an Engineered Microbiome"

### SUPPLEMENTARY MATERIALS AND METHODS

#### Metagenomic Library Construction and Sequencing

Samples were collected from a single bioreactor simulating enhanced biological phosphorus removal as described previously (1, 2). Briefly, a laboratory-scale sequencing batch reactor (SBR) with a 2L working volume was seeded with activated sludge from the Nine Springs WWTP located in Madison, WI, USA. The reactor was fed primarily with acetate, and operated in four daily cycles of 6 hours, with a hydraulic residence time (HRT) of 12 hours, and a solids retention time (SRT) of four days. The total anaerobic/aerobic cycle time was 6 hours with 140 min anaerobic contact (sparging with $N_2$ gas) and 190 min aerobic contact (sparging with air), followed by a 30 min settling period. Biomass samples for metagenomic sequencing were collected by centrifuging 2 mL of mixed liquor at 8,000 *x g* for 2 min, and DNA was extracted using a modified phenol:chloroform bead-beating extraction protocol.

100 ng of DNA was sheared to 300 bp using the Covaris LE220 (Covaris) and size selected with SPRI beads (Beckman Coulter). The fragments were ligated with end repair, A-tailing, Illumina compatible adapters (IDT Inc) using the KAPA-Illumina library preparation kit (KAPA Biosystems). Libraries were quantified using the KAPA Biosystem next-generation sequencing library qPCR kit and ran on a Roche LightCycler 480 real-time PCR instrument. The quantified libraries were prepared for sequencing on the Illumina HiSeq platform using the v4 TruSeq paired-end cluster kit and the Illumina cBot

instrument to create a clustered flow cell for sequencing. Shotgun sequencing was performed for ten samples at the Department of Energy Joint Genome Institute (Walnut Creek, CA, USA) on the Illumina HiSeq 2500 platform with the TruSeq SBS sequencing kits, followed by 2x150 indexing. All metagenomic libraries consist of ~100 million 150-bp Illumina HiSeq reads with approximately 15 Gb per sample.

**Metagenomic Assembly, Annotation, and Metatranscriptomic Mapping**

Raw metagenomic reads were quality filtered and trimmed using bbduk as part of the BBtools suite v38.07 (3). Each of the three metagenomic samples were individually assembled into contigs using metaSPAdes (4). Additionally, all three metagenomes were co-assembled using metaSPAdes and MEGAHIT (4, 5). Metagenomic reads from each time-point were mapped against all assemblies using BBMap with a 95% sequence identity cutoff. Assembled contigs from each time-point were binned into population genomes using MetaBAT2 v2.12.1 informed by the differential coverage of all time-points (6). To determine the most high-quality set of non-redundant bins across time-points and assembly methods, we calculated pairwise ANI for each set of bins across all time-points and dereplicated with dRep v2.4.2 (7). Of the 66 genomes, 15 are high quality draft genomes, with >95% completeness, <5% redundancy, contain all 3 ribosomal RNAs (5S, 16S, and 23S) and more than 18 tRNAs, according to MiMAGs standards (8). No two sets of bins are similar by more than 85% ANI, reflecting the well-established sequence-based species cutoff (9, 10). Each genome was manually inspected for uniform coverage across all contigs using Anvi'o v5.0 (11). Draft genomes were classified using both the GTDB-tk v0.13 automatic classification method (12, 13) and a manually curated taxonomy assignment based on concatenated ribosomal protein phylogenies (14, 15). A phylogenetic tree of concatenated ribosomal proteins of all 66 bins, select reference genomes, and a subset of high-quality genomes assembled from Danish WWTPs described in Singleton et al. (16), was constructed using the metabolisHMM package with tree building performed by RaxML and visualized using iTOL (15, 17, 18) (Supplementary Figure 1) iTOL tree is available online at https://itol.embl.de/tree/978318524309771596154125#.

### SUPPLEMENTARY FIGURES AND TABLES

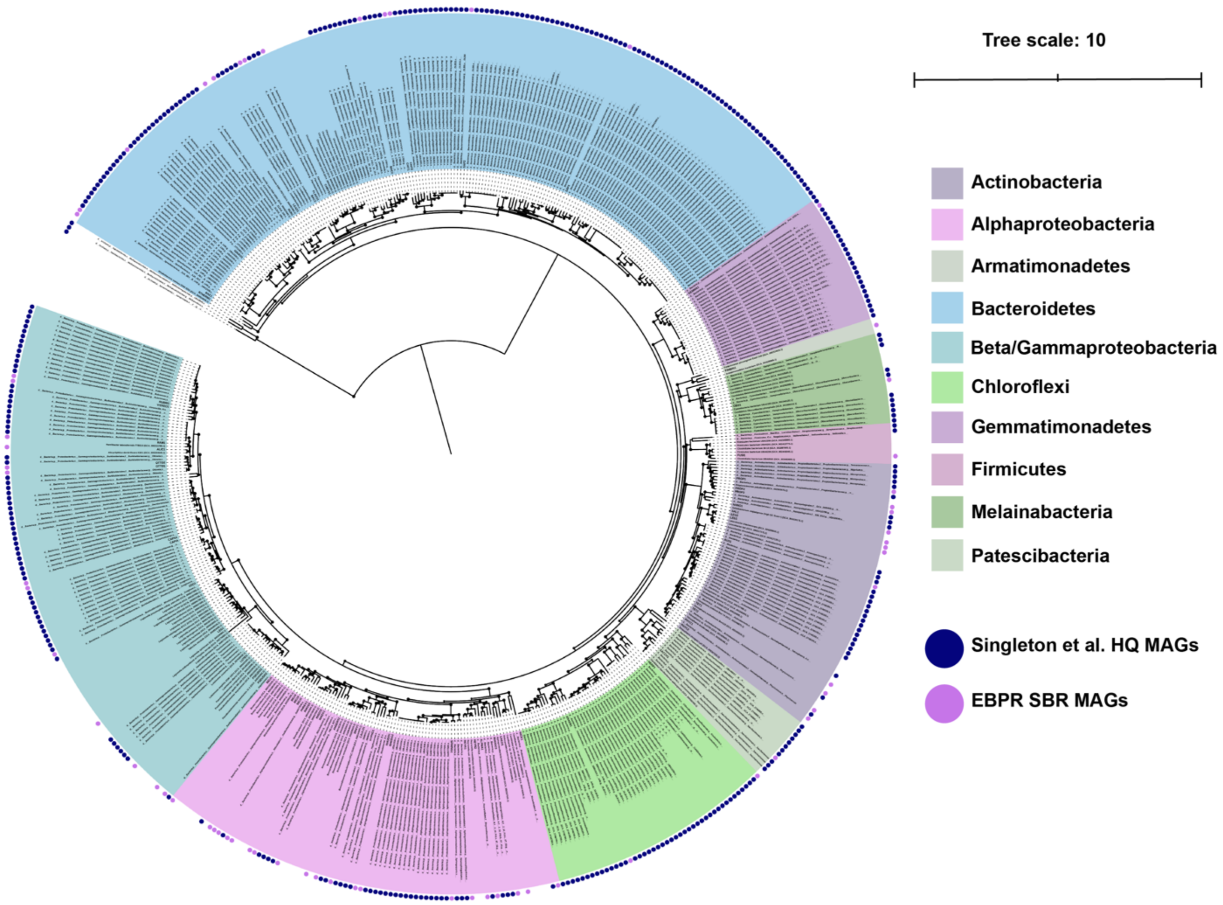


**Supplementary Figure 1.** Phylogenetic tree of all 66 assembled MAGs compared to a subset of high-quality MAGs assembled by Singleton et al. from a full-scale WWTP and select references from Genbank. The tree was constructed with concatenated single marker genes from the GTDB-tk separately for bacterial and archaeal genomes. The two alignments were combined with muscle and the tree was constructed with RaxML using automatic bootstrap support. The tree was visualized and annotated using iTOL and can be viewed interactively at https://itol.embl.de/tree/978318524309771596154125#.

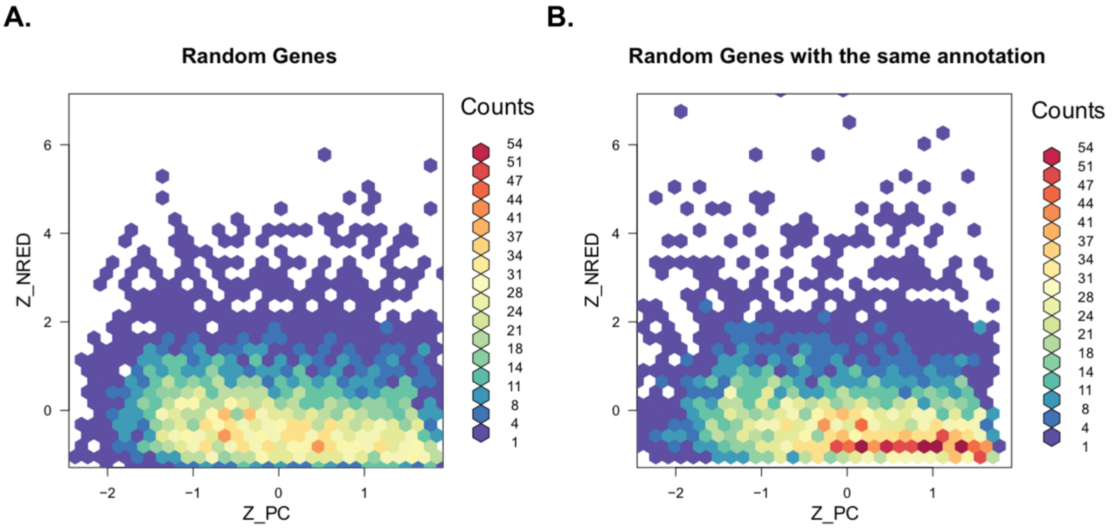


**Supplementary Figure 2.** Benchmarking of the random background distributions of distance between genes. Two random background distributions were benchmarked by calculating the distance (Pearson Correlation and Normalized Rank Euclidean Distance) between randomly sampled genes. A) Completely random pairs of genes were sampled. B) Pairs of genes were sampled that share the same randomly picked KEGG Orthology term.

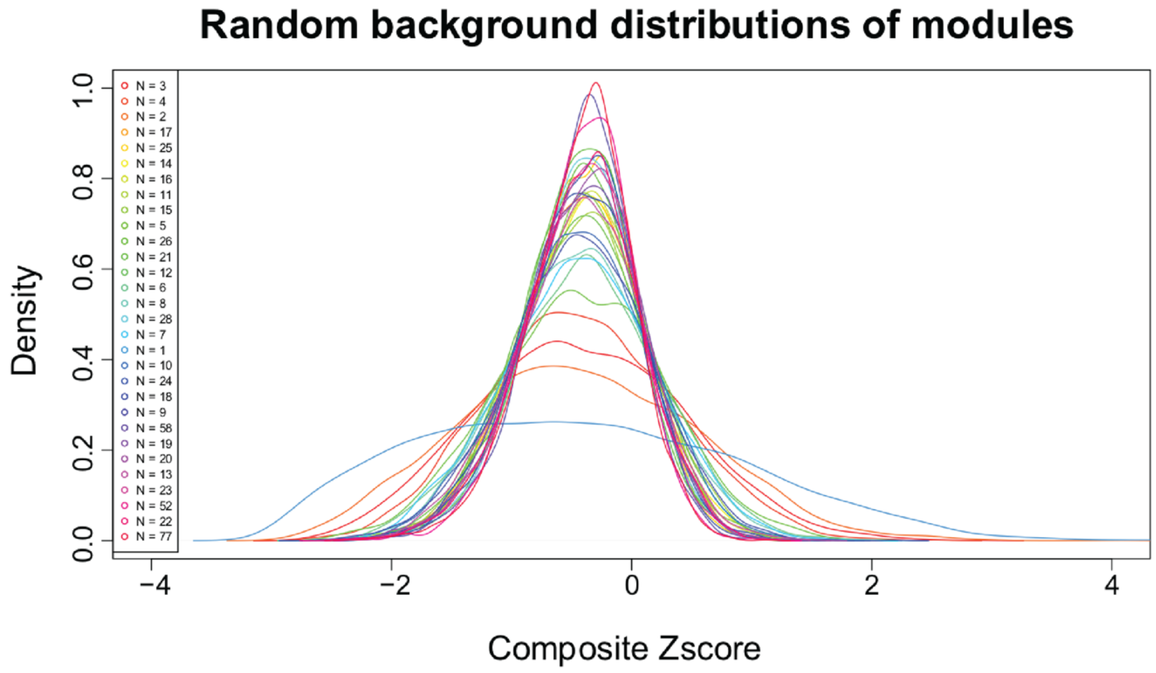


**Supplementary Figure 3.** Benchmarking of the random background distributions of distances between traits. Random traits were sampled containing N number of genes (N is equivalent to the possible number of genes in the trait library). The distances between two traits were calculated as a composite Z score of the Pearson Correlation and the Normalized Rank Euclidean Distance.

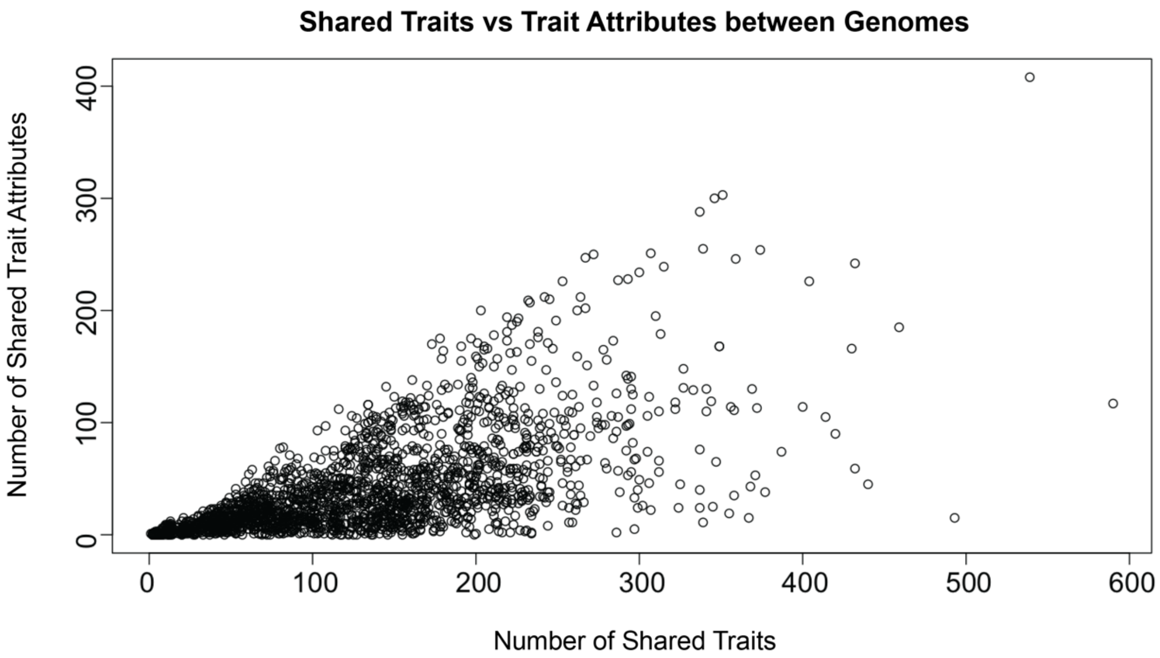


**Supplementary Figure 4.** Benchmarking of the number of traits versus the number of trait-attributes that pairs of genomes share. For each combination for genomes, the number of shared traits, and the number of shared trait-attributes were calculated.

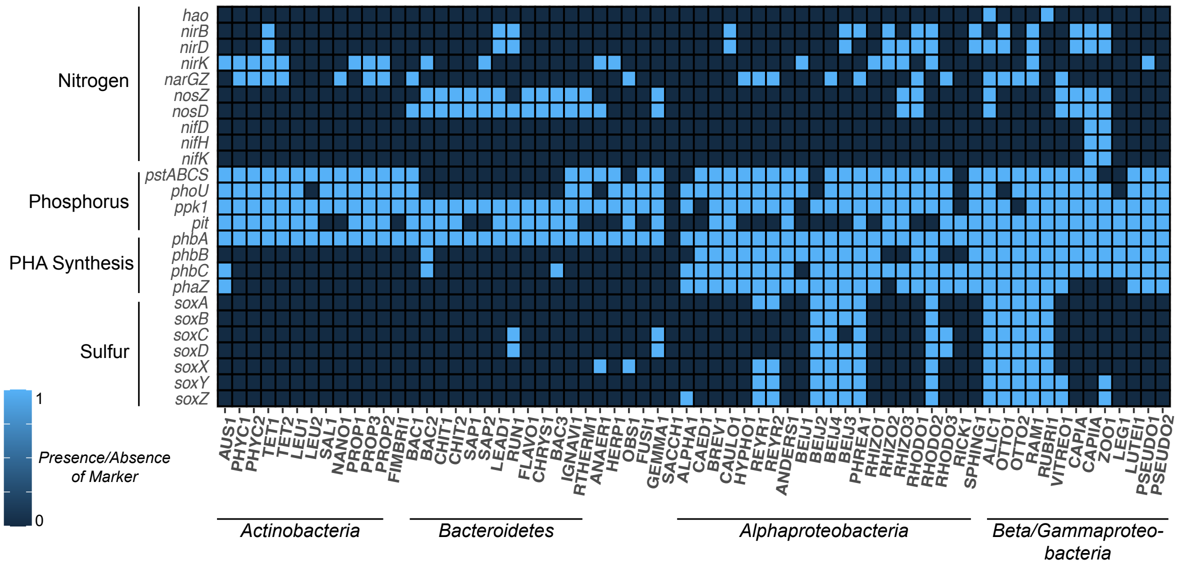


**Supplementary Figure 5. Presence of EBPR-related Traits**

Presence-absence heatmap of select genes underlying key EBPR traits such as nitrogen cycling, phosphorus transport, PHA synthesis, and sulfur oxidation.

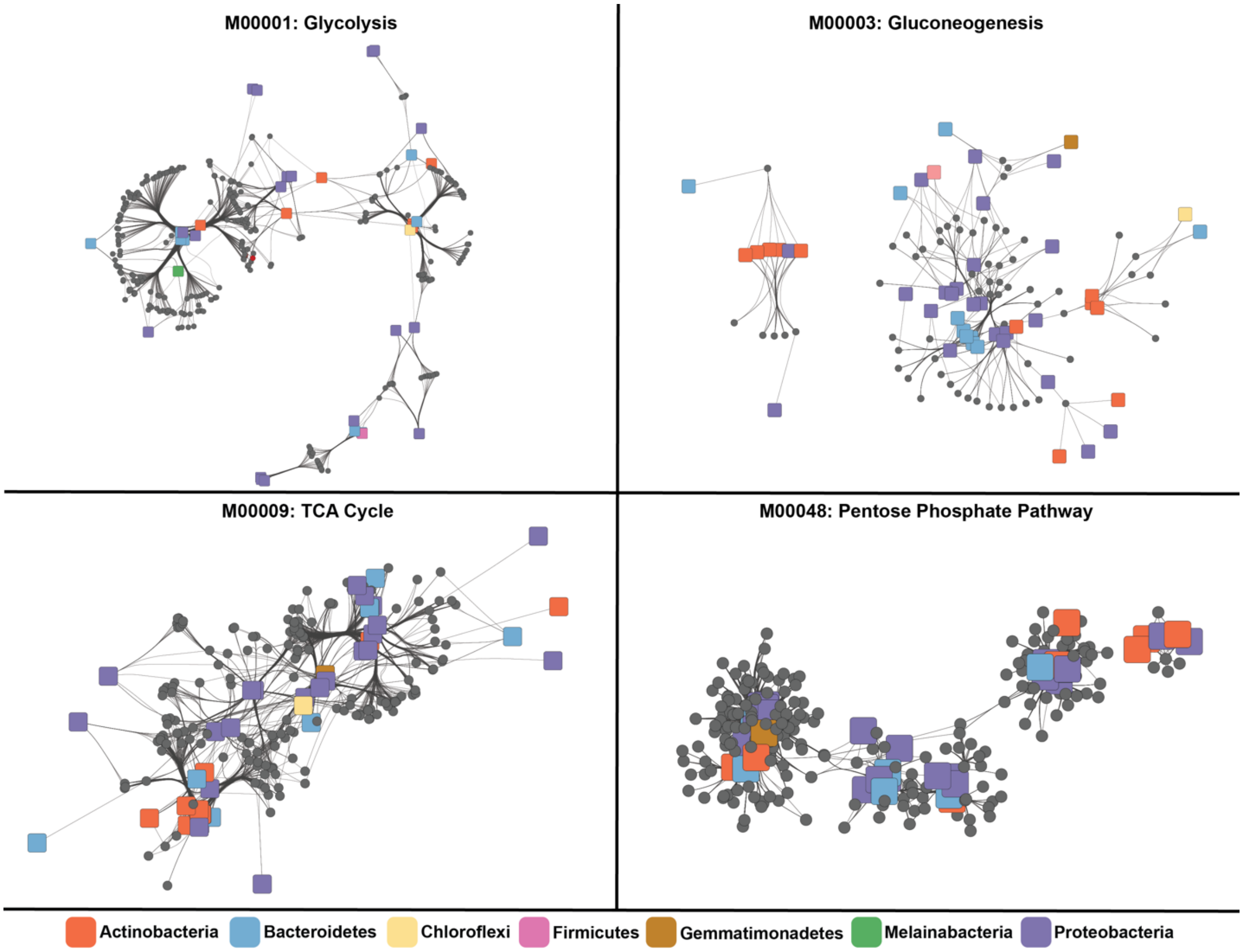


**Supplementary Figure 6. Trait-Attributes of Large Modules**

Individual networks of large KEGG modules with multiple routes. Squares represent individual genomes colored by phylum, and nodes are different parts of the module

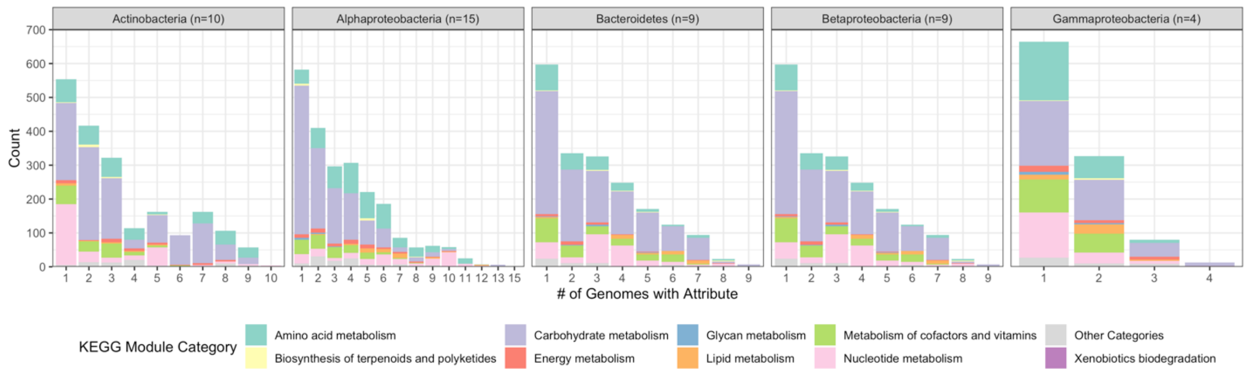


**Supplementary Figure 7.**

Expansion of Figure 3B summarized by phylum. For each phylum the total number of genomes is given, with each bar representing the number of trait attributes present with that number of genomes within the phylum. Stacked bars are colored by KEGG module category. Tables for core (present in greater than more than half of the genomes within each phylum) and niche differentiating (present in less than half of the genomes within each phylum) for each phylum are available in the Extended Table 1 available on Figshare https://figshare.com/articles/dataset/Lineage-Specific_Core_and_Niche_Differentiating_Traits/15001200.

| Max samples zero counts | Genes | Significant Trait Attributes |
|---|---|---|
| 0 | 97500 | 1938 |
| 1 | 123423 | 1990 |
| 2 | 142426 | 1687 |
| 3 | 159843 | 1873 |
| 4 | 178446 | 1674 |

**Supplementary Table 1.** Benchmarking results of the number of samples in the time-series allowed to have zero RNAseq counts per individual gene. The benchmarks range from strict, where zero samples are allowed to have an expression count of zero for each gene, to loose, where four samples are allowed to have a count of zero for each gene. The genes that do pass the threshold value of N samples were pruned from the table. For each benchmarking iteration, the number of genes that remained was recorded, together with the total number of statistically significant trait attributes.
